# Intra-amygdala circuits of sleep disruption-induced anxiety in female mice

**DOI:** 10.1101/2024.05.19.594863

**Authors:** Beibei Peng, Allison R. Foilb, Yunona Manasian, Yan Li, Xin Deng, Edward G. Meloni, Kerry J. Ressler, William A. Carlezon, Vadim Y. Bolshakov

## Abstract

Combining mouse genetics, electrophysiology, and behavioral training and testing, we explored how sleep disruption may affect the function of anxiety-controlling circuits, focusing on projections from the basolateral nucleus of the amygdala (BLA) to CRF-positive cells in the lateral division of the central amygdala (CeL). We found in Crh-IRES-Cre::Ai14(tdTomato) reporter female mice that 6 hours of sleep disruption during their non-active (light) cycle may be anxiogenic. Notably, the AMPAR/NMDAR EPSC amplitude ratio at the BLA inputs to CRF-CeL cells (CRF^CeL^), assessed with whole-cell recordings in *ex vivo* experiments, was enhanced in slices from sleep-disrupted mice, whereas paired-pulse ratio (PPR) of the EPSCs induced by two closely spaced presynaptic stimuli remained unchanged. These findings indicate that sleep disruption-associated synaptic enhancements in glutamatergic projections from the BLA to CRF-CeL neurons may be postsynaptically expressed. We found also that the excitation/inhibition (E/I) ratio in the BLA to CRF^CeL^ inputs was increased in sleep-disrupted mice, suggesting that the functional efficiency of excitation in BLA inputs to CRF^CeL^ cells has increased following sleep disruption, thus resulting in their enhanced activation. The latter could be translated into enhanced anxiogenesis as activation of CRF cells in the CeL was shown to promote anxiety-like behaviors.

## Introduction

Sleep disruption, including reduced sleep duration and quality, is highly prevalent in human subjects’ life, imposing serious social burdens (Hillman and Lack, 2013; Madan and Jha, 2023). Chronic sleep loss has deleterious consequences for the central nervous system function, immune reactivity, and cardiovascular health, ultimately resulting in severe physical and mental dysfunctions (Palma et al., 2013; Besedovsky et al., 2019; Khan and Aouad, 2022). Notably, female subjects are more vulnerable to sleep loss-associated impairments compared to males, exhibiting higher susceptibility to the mentioned deleterious outcomes (Paul et al., 2006; Zhang and Wing, 2006; Hajali et al., 2012; Hillman and Lack, 2013; Foilb et al., 2023). The life-time rate of developing an anxiety disorder in women is approximately two times higher than in men, and this disease is often associated with sleep disturbances (Kessler et al., 1994; Ramsawh et al., 2009; Altemus et al., 2014; Cox and Olatunji, 2016). The relationship between sleep loss and anxiety disorders appears to be bidirectional and reciprocal (Alvaro et al., 2013), with anxiety preceding and predicting sleep disturbances (Ohayon and Roth, 2003; Johnson et al., 2006), whereas sleep disturbances exacerbate anxiety symptom severity (Selvi et al., 2007; Babson et al., 2010; Cox and Olatunji, 2016; Pires et al., 2016a). Furthermore, female subjects are more vulnerable to developing anxiety symptoms after sleep deprivation than males (Goldstein-Piekarski et al., 2018). Animal studies, however, have shown inconsistent results in respect to sleep disruption-associated changes in anxiety levels (Pires et al., 2015).

The amygdala, a brain structure located in the medial temporal lobe, is thought to play pivotal roles in the mechanisms of fear control (Davis, 1992; Sah et al., 2003; Shin et al., 2006; Yoo et al., 2007; van der Helm et al., 2011). Whereas the amygdala is structurally heterogenous, its two specific subdivisions—basolateral amygdala (BLA) and lateral subnucleus of the central amygdala (CeL)—were repeatedly shown to play essential functions in innate fear (anxiety) (Sanders and Shekhar, 1995; Tye et al., 2011; Ahrens et al., 2018), and recent studies demonstrated that these two sites are also involved in sleep regulation (Ma et al., 2019; Montes-Rodríguez et al., 2019; Wang et al., 2020; Hasegawa et al., 2022). Interestingly, CeL is densely populated by corticotropin-releasing factor (CRF)-expressing neurons (CRF^CeL^) which receive strong excitatory inputs from BLA and contribute to both anxiety and sleep mechanisms (Fadda and Fratta, 1997; Pawlyk et al., 2006; Sanford et al., 2017; McCullough et al., 2018, 2020; Pomrenze et al., 2019b, 2019a; Jo et al., 2020), suggesting that sleep disruption-associated changes in neurotransmission in BLA-to-CRF^CeL^ projections may contribute to sleep disruption triggered anxiety enhancements. Here we addressed this possibility at synaptic and neural circuit-function levels.

## Methods

### Animals

CRF reporter mice were generated by crossing CRH^IRES-CRE^ mice with Ai14 (tdTomato) mice. Adult female mice (2-4 months old) were used in this study. Animals were housed in groups in a temperature-controlled room with 12:12-h light:dark cycle, with light on at 7:00 A.M. The light cycle (non-active cycle) started at 7:00 AM and lasted until 7:00 PM, whereas the dark cycle (active cycle) started at 7:00 PM and lasted until 7:00 AM. Mice in the sleep disruption group (SD) were subjected to the sleep disruption procedures for 6 hours from 7:00 AM to 1:00 PM (during non-active cycle) in the sleep fragmentation chamber (Lafayette Instrument Company, Model 80391). Control (CON) mice were subjected to the same procedures for 6 hours from 7:00 PM to 1:00 AM (in their awake cycle). Mice from both groups were tested behaviorally or electrophysiologically at the same time, starting at 1:00 PM. Food and water were provided ad libitum. All animal care and experimental procedures were approved by the McLean Hospital Institutional Animal Care and Use Committee.

### Histology and microscopy

Brain slices containing biocytin injected cells were fixed in 4 % paraformaldehyde in PBS at 4°C overnight, followed by washing in PBS three times, for 5 min each time, and then incubated in 0.1% Triton PBS for an hour at room temperature. Sections were then incubated in 0.4% Triton PBS containing streptavidin Alexa 488 conjugate (10 μg/ml) overnight at room temperature. After that, sections were washed three time in PBS followed by Hoechst (1μg/ml) staining for 10 min. Leica TCS SP8 confocal microscope fluorescent microscope was used to capture images which were processed with ImageJ software (NIH).

### In vitro electrophysiological recordings

Coronal brain slices (300 µm in thickness) containing BLA and CeL were prepared with a vibratome in an ice-cold solution containing (in mM): 252 sucrose, 2.5 KCl, 5.0 MgCl_2_, 1.0 CaCl_2_, 1.25 NaH_2_PO_4_, 26 NaHCO_3_, 10 glucose, equilibrated with 95% O_2_ and 5% CO_2_. Slice were then incubated into oxygenated artificial cerebrospinal fluid (ACSF) containing (in mM): 125 NaCl, 2.5 KCl, 2.5 CaCl_2_, 1.0 MgSO_4_, 1.25 NaH_2_PO_4_, 26 NaHCO_3_ and 10 glucose for 30 min at 32°C and then at room temperature. CRF positive neurons in CeL were targeted for patch-clamp recordings using the fluorescent microscope (Zeiss, Examiner.A1). In the experiments in current-clamp recording mode, recording patch electrodes (3-5 MΩ resistance) were filled with the with potassium gluconate-based internal solution containing (in mM): 130 K-gluconate, 5.0 KCl, 2.5 NaCl, 1.0 MgCl_2_, 0.2 EGTA, 10 HEPES, 2.0 MgATP, 6.0 neurobiotin and 0.1 NaGTP (adjusted to pH 7.2 with KOH). The 500-ms-long current injections, delivered through the recording electrode and ranging from 0 to 200 pA (with the 25 pA increment), were used to assess neuronal excitability. For action potential waveform analysis, the first, middle and last action potentials were extracted from responses to 100 pA-currents injected into the recorded neuron. The action potential threshold was determined as the voltage at which the rising slope was approaching 10 V/s (Brownstone et al., 1992). I_h_ currents were recorded using potassium gluconate-based internal solutions in voltage-clamp mode delivering 500-ms voltage-steps ranging from -60 to -130 mV (with the 10 mV increment) to the recorded neuron. Ih currents were obtained by calculating the difference between the initial phase and the steady-state phase after the voltage injection (Baimel et al., 2017). In other voltage-clamp recording, Cs-methanesulfonate-based internal solution was used, containing (in mM): 135 CsMeSO_4_, 5.0 NaCl, 1.0 MgCl_2_, 0.2 EGTA, 10 HEPES, 6.0 neurobiotin, 2.0 ATP and 0.2 GTP. Whole-cell recordings were performed at 30-32 °C using EPC-9 amplifier and Pulse 8.8 software (Heka Elektronik). Currents were filtered at 1 kHz and digitized at 5 kHz. Silver painted patch pipettes filled with ACSF were positioned in the BLA to evoke synaptic currents in CRF^CeL^ neurons. Inhibitory postsynaptic currents (IPSCs) were blocked by GABA_A_ receptor antagonist bicuculline (BIC, 10 μM, R&D systems) in the course of sEPSCs recordings, as well as in the experiments assaying paired-pulse ratio (PPR) and the AMPAR/NMDAR EPSC amplitude ratio. NBQX (10 μM, R&D systems) was used to block glutamatergic inputs to CRF^CeL^ neurons. To obtain estimates of the AMPAR/NMDAR EPSC amplitude ratio, the NMDAR- and AMPAR-mediated EPSCs were recorded at +40 and -70 mV, respectively. The amplitude of AMPAR EPSCs was measured at the peak at -70 mV, and the amplitude of NMDAR EPSCs was measured at +40 mV at 40 ms after the peak at -70 mV. PPR was induced by two closely spaced presynaptic stimuli with 50 ms inter-stimulus interval at -70 mV. To obtain the excitation/inhibition ratio (E/I ratio) estimates, the monosynaptic EPSCs were recorded in CRF^CeL^ neurons at –70 mV (close to the reversal potential for the IPSC) first, and then we recorded IPSCs at 0 mV (the reversal potential for AMPAR EPSCs) in same neurons.

### Behavioral testing

Mice were transferred to the behavioral testing room and acclimated in the room for 30 min prior to behavioral testing. The elevated plus-maze was made of plastic and consisted of two open arms (30×5 cm) and two enclosed arms (30×5 cm) extending from a central platform (5 x 5 x 5 cm) at 90 degrees. The maze was placed 80 cm above the floor. Red light (10 lux) was used in the EPM tests. At the start of each session, a mouse was placed in the center of the EPM facing one of the closed arms. Mice explored the EPM apparatus for 10 min and behavior was videotaped for the subsequent analysis. Four hours after the EPM test, mice were subjected to the open field test. Mice were placed in one corner of a brightly lit (100 lux) open field, which was made of white plastic board in size of 45 × 45 × 30 cm (L× W× H). Mice explored the open field for 10 min. EthoVision software was used to capture and analyze the results.

### Statistical analysis

For statistical comparisons, we used unpaired Student’s t-test or ANOVA as indicated in the description of specific experiments. Statistical analysis was performed using Prism 10 (GraphPad Prism) software. P < 0.05 was considered statistically significant.

## Results

### Acute sleep disruption results in enhanced anxiety-like behaviors in female mice

To explore the circuit level mechanisms of sleep disruption-associated behavioral impairments, we generated CRF reporter mice (see **Methods)**. These mice, expressing tdTomato in CRF-positive neurons, were used in both behavioral studies and *ex vivo* electrophysiological experiments.

It has been found previously that 6 hours of total sleep deprivation by gentle handling were sufficient to produce anxiogenic effects in rats as assayed with grooming analysis algorithm (Pires et al., 2013). Moreover, sleep disruption with the sweeper bar in the sleep fragmentation chamber in mice was shown to be anxiogenic in the elevated plus-maze test (Nair et al., 2011), a standard behavioral assay quantifying anxiety-like behaviors in mice (Bourin et al., 2001). Based on these earlier studies, we designed the sleep disruption protocol, placing mice from the SD group to the sleep fragmentation chamber (Lafayette Instrument Company, Model 80391) for 6 hours during inactive cycle (light cycle; see **Methods** for details). Mice sleep 2-4 times more during inactive cycle (light cycle) than during active phase (dark cycle) (Koehl et al., 2003). As 6-hr-long sleep disruption with the sweeper bar during light cycle might be stressful for mice (Foilb et al., 2024), our control group of mice was subjected to the same sweeper bar disturbances in the sleep fragmentation chamber but in the early dark (active) phase which did not interfere with sleep (Bian et al., 2022) (Fig. 1A). Thus, we controlled for the functional effects not directly related to sleep disruption. After sleep disruption procedures, mice were subjected to elevated plus-maze (EPM) or open field (OF) tests (Fig. 1A,B), which are frequently used to assay anxiety-like behaviors (Riccio et al., 2009; Kim et al., 2013). We found that mice from SD group spent significantly less time in the open arms of the EPM and entered them less frequently compared to CON mice (unpaired *t*-test; for number of entries: *t*_(16)_ = 2.207, *p* = 0.042, for open arm time: *t*_(16)_ = 3.444, *p* = 0.003 between SD and CON groups (Fig. 1C, E). There was no difference in the travel distance between the groups (unpaired *t*-test; *t*_(16)_ = 1.007, *p* = 0.3291) (Fig. 1C, E), indicating that our SD procedures did not induce fatigue or mania-like hyperactivity (Wu et al., 2023). In the open field test (OF), while the differences in the number of entries to the OF center did not reach the statistical significance level (unpaired *t*-test; *t*_(16)_ = 1.627, *p* = 0.123), the center area time was significantly decreased in SD group compared to CON mice (unpaired *t*-test; *t*_(16)_ = 2.429, *p* = 0.027) (Fig. 1D, F). Similarly to EPM results, the total travel distance during OF test was not different between CON and SD mice (unpaired *t*-test; *t*_(16)_ = 1.721, *p* = 0.105) (Fig. 1D, F). These findings indicate that SD during non-active (light) phase may be associated with enhanced anxiety levels in female mice.

**Figure 1.**
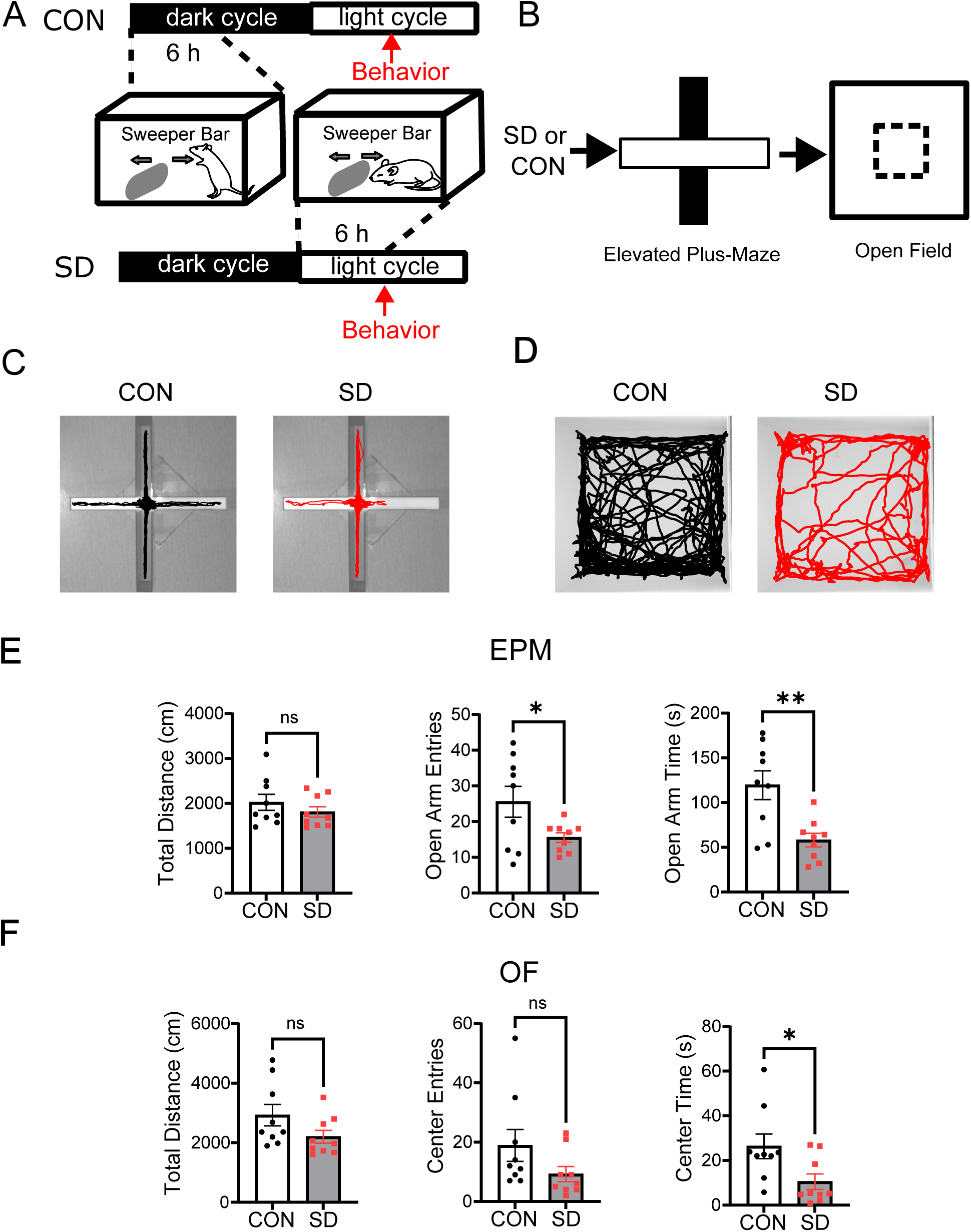
Sleep disruption enhances anxiety-like behavior in female mice. **A**, A diagram showing how the experiments were designed. CON or SD mice were placed into the sleep fragmentation apparatus (see **Methods**) during the early dark or light phases of the light:dark cycle, respectively. After SD, both groups were subjected to the behavioral tests. **B**, Timeline of behavioral testing. Mice were tested in EPM first, and it was followed by OF with the interval of four hours between these two tests. **C**, Representative behavioral tracks in EPM test. **D**, Representative behavioral tracks in OF test. **E**, The total distance (left) was unchanged after SD. SD exhibited decreased time in the open arms (middle) and the number of entries to the open arms (right) in EPM test; n = 9 mice per each group. **F**, The total distance (left) and center entries (middle) were unaffected after SD in OF test. SD mice exhibited decreased time spent in the center (right); n = 9 mice per each group.

### Sleep disruption does not affect membrane properties of CRF^CeL^ neurons

It has been repeatedly shown previously that BLA and CeL are the key brain structures controlling the mechanisms of anxiogenesis (Tye et al., 2011; Felix-Ortiz et al., 2013; Cai et al., 2014; Pomrenze et al., 2019b, 2019a; Douceau et al., 2022). Thus, we hypothesized that SD-associated plastic changes in BLA-CeL circuits could contribute to SD-triggered anxiety enhancements (Fig. 1C-F). As activation of CRF^CeL^ neurons was shown to promote anxiety-like behaviors (Pomrenze et al., 2019b, 2019a), we first addressed the possibility that CRF^CeL^ cells excitability might be affected in sleep disrupted mice. For this, we performed whole-cell recordings from CRF^CeL^ neurons in slices from CRF-reporter mice from SD or control groups of mice which were subjected to the identical (time-matched) SD or CON procedures, respectively, as in the experiments probing the effects of SD on anxiety levels (Fig. 2A-C). We found that the number of action potentials triggered by prolonged depolarizing current injections of increasing intensity under current-clamp conditions was not different between CON and SD mice (2-way ANOVA, F_(1,37)_ = 0.385, *p* = 0.539; Fig. 2D), indicating that acute SD did not affect membrane excitability of CRF^CeL^ neurons. Moreover, the resting membrane potential was unaffected in SD mice (unpaired *t*-test, *t*_(37)_ = 1.092, *p* = 0.282) (Fig. 3A). The waveform of action potentials also remained unchanged in sleep disrupted mice (Fig. 3B), as there were no differences between CON and SD groups in the action potential amplitude (two-way ANOVA; F_(1,30)_ = 0.090, p = 0.766; Fig. 3C), AP threshold (F_(1,30)_ = 1.128, *p* = 0.297; Fig. 3D), afterhyperpolarization (F_(1,30)_ = 1.368, *p* = 0.251; Fig. 3E), maximum rising phase slope (F_(1,30)_ = 0.105, *p* = 0.748) (Fig. 3F), falling phase slope (F_(1,30)_ = 0.259, *p* = 0.615; Fig. 3G) and width (F_(1,30)_ = 0.081, *p* = 0.779; Fig. 3H). Hyperpolarization-activated cyclic nucleotide-gated (HCN) channels-mediated currents (Ih), triggered by delivering hyperpolarizing voltage steps into recorded CRF^CeL^ neurons, were also not different between the groups (F_(1,33)_ = 1.594, p = 0.216) (Fig. 3I,J). These results indicate that SD had no effect on the membrane properties of CRF^CeL^ neurons.

**Figure 2.**
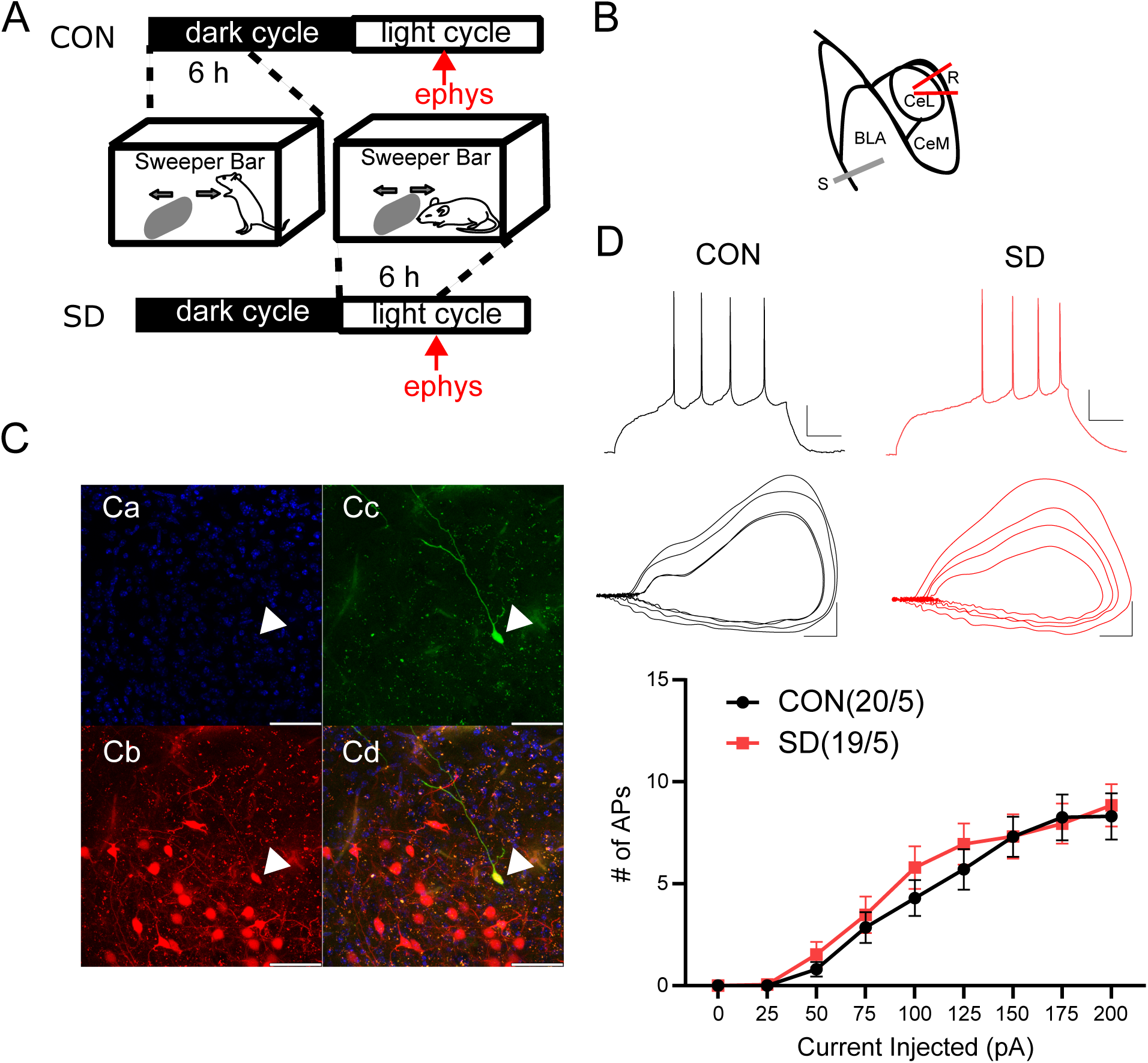
SD had no effect on membrane excitability of CRF^CeL^ neurons. **A**, A diagram of experimental design of electrophysiological studies. Mice were sacrificed for electrophysiological experiments at times matching behavioral testing. **B**, CRF reporter mice were used to record from the CRF positive neurons in the CeL. To activate the BLA to CRF^CeL^ projections, a stimulating electrode was placed in the BLA. **C,** panel **Ca** shows Hoechst staining of the brain section containing CeL from a CRF-reporter mouse. **Cb**, Microscopic mage of the same brain section showing CRF positive neurons. **Cc**, the neurobiotin-filled recorded cell that was CRF positive. **Cd**, the overlayed **Cb** and **Cc** images. Scale bar, 50 µm. **D**, Membrane excitability of CRF^CeL^ neurons was unchanged after SD. Top: representative traces action potentials induced by depolarizing current injections (75 pA). Scale bars: 20 mV, 100 ms. Middle: Representative traces of phase plots of action potentials induced by depolarizing current injections. Scale: 50 V/s, 10 mV. Lower, Membrane excitability was unchanged in SD mice, CON: n = 20 neurons from 5 mice; SD: n = 19 neurons from 5 mice.

**Figure 3.**
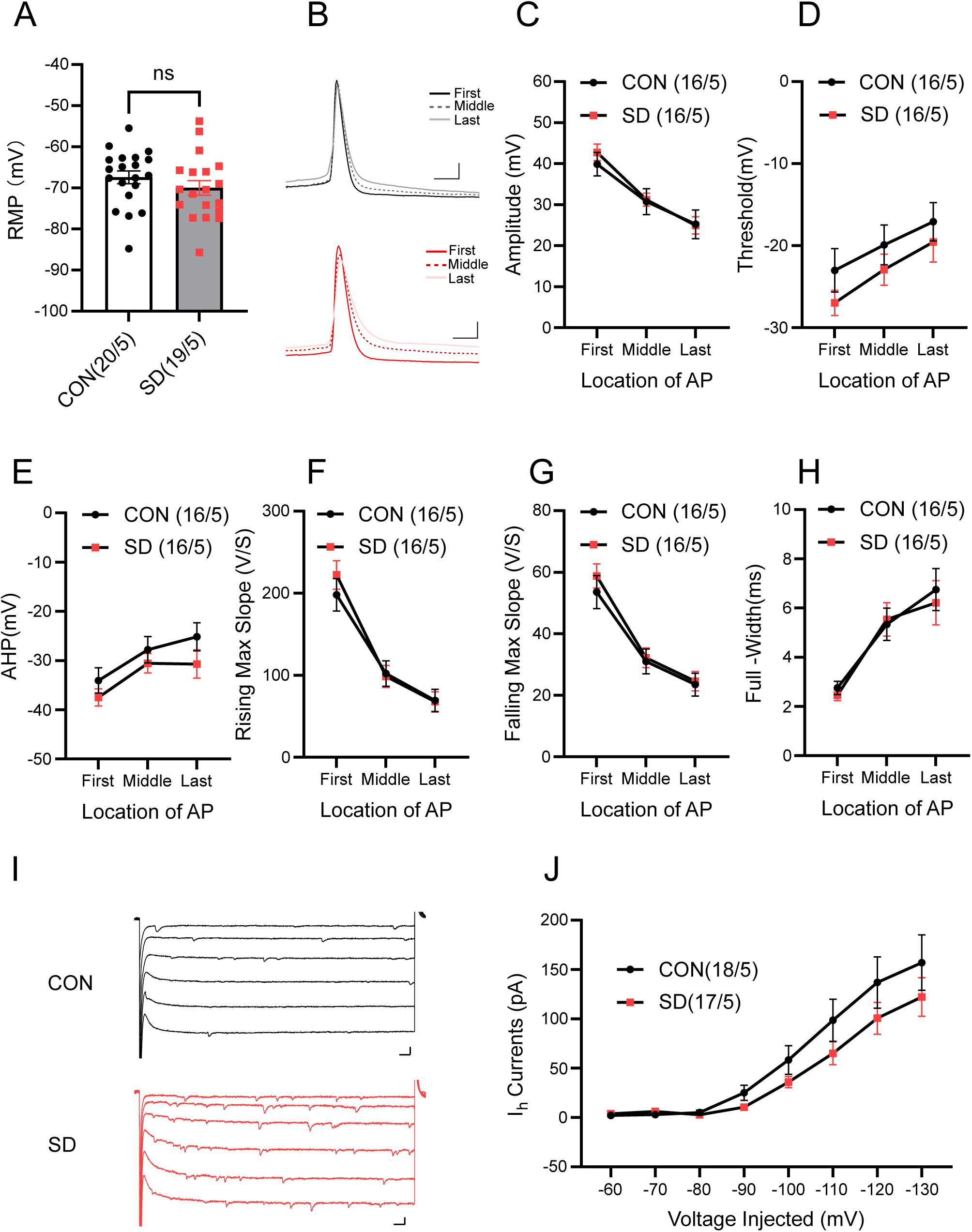
Intrinsic membrane properties of CRF^CeL^ neurons were unaffected in SD mice. A, Resting membrane potential (RMP) of CRF^CeL^ neurons was unchanged after S; CON, n = 20 neurons from 5 mice; SD, n = 19 neurons from 5 mice. **B**, Representative traces of step-current evoked action potentials. Waveforms of first, middle and last action potential during one depolarizing step in CON mice (top, black) and SD mice (lower, red). Scale: 10 mV, 2.5 ms. **C**, Amplitude of current injection-evoked action potential was not different between CON and SD mice. **D**, Threshold of action potential generation was unaffected in SD mice. **E**, Afterhyperpolarization (AHP) of action potentials was unchanged in SD mice. **F**, Maximum slope of the rising phase of the action potential was unaffected in SD mice. **G**, Maximum slope of the falling phase of the action potential was unaffected in SD mice. **H**, Width of action potentials was unchanged in SD mice. **C-H**, CON, n = 16 neurons from 5 mice; SD, n = 16 neurons from 5 mice. I, Representative I_h_ currents in CRF^CeL^ neurons in slices from CON (black) and SD (red) mice, triggered by voltage steps from -80 to -130 mV. Scale, 20 pA, 20 ms. J, I_h_ currents were unchanged after SD. CON, 18 neurons from 5 mice; SD, 17 neurons from 5 mice.

### Spontaneous excitatory and inhibitory synaptic drive at inputs to CRF^CeL^ neurons is enhanced and decreased, respectively, in sleep-disrupted female mice

As there were no alterations in intrinsic membrane properties of CRF^CeL^ neurons in SD mice, we next tested a possibility that the function of local neuronal networks in the CeL might be affected in sleep disrupted mice, modifying spontaneous synaptic drive at inputs to CRF-positive cells. Thus, we first recorded spontaneous excitatory postsynaptic currents (sEPSCs) in CRF^CeL^ neurons at a holding potential of -70 mV under voltage-clamp conditions. We found that both the frequency (unpaired *t*-test; *t*_(17)_=2.700, *p* = 0.015) and amplitude (*t*_(17)_=2.457, *p* = 0.025) of sEPSCs were increased in CRF positive cells in slices from SD mice (Fig. 4 A-C). Unexpectedly, the frequency of spontaneous inhibitory postsynaptic currents (sIPSCs), recorded at a holding potential of 0 mV under voltage-clamp conditions and mediated by GABA_A_ receptors, in CRF^CeL^ neurons was decreased in SD group compared to CON mice (unpaired *t*-test; *t_(_*_29)_=3.312, *p* = 0.003) (Fig. 4D,E), whereas the amplitude of sIPSCs remained unchanged (*t*_(29)_=1.788, *p* = 0.084) (Fig. 4D,F). The observed enhancements of excitatory and diminished inhibitory drive indicate that the overall level of tonic excitation of CRF^CeL^ neurons (by sEPSCs) may be enhanced in sleep disrupted mice, thus possibly augmenting synapticly-driven spiking in specific pathways.

**Figure 4.**
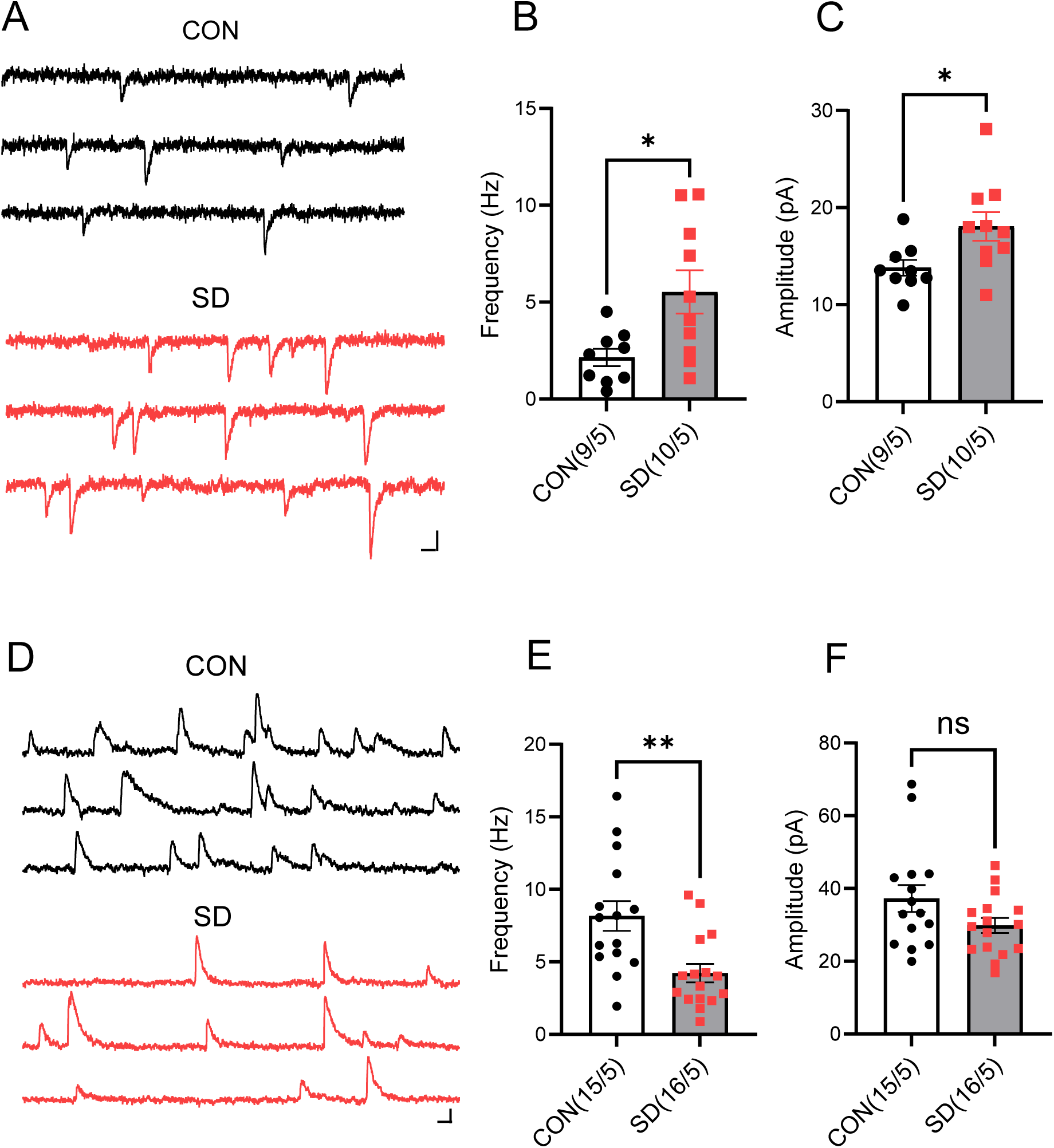
Spontaneous synaptic excitatory and inhibitory drives to CRF^CeL^ neurons in SD mice. **A**, Representative traces of sEPSCs in CRF^CeL^ neurons. Scale bars: 10 pA, 10 ms. **B**, Sleep disruption resulted in enhanced frequency of sEPSCs in CRF^CeL^ neurons. **C**, The sEPSC amplitude was increased after SD. **B-C**, CON, 9 neurons from 5 mice; SD, 10 neurons from 5 mice. The sEPSCs were recorded at a holding potential of -70 mV. **D**, Representative traces of sIPSCs. Scale: 20 pA, 10 ms. D, Frequency of sIPSCs in CRF^CeL^ neurons was decreased in SD mice. **F**, Amplitude of sIPSCs was unchanged after SD. **E-F,** CON. 15 neurons from 5 mice; SD, 16 neurons from 5 mice.

### Synaptic efficacy in BLA projections to CRF^CeL^ neurons is enhanced in SD mice

Consistent with our earlier results (Cho et al., 2012), electrical stimulation of BLA neurons resulted in biphasic synaptic responses consisting of the monosynaptic glutamatergic excitatory postsynaptic current (EPSC) and delayed inhibitory postsynaptic current (IPSC) in CRF^CeL^ neurons (Fig. 5A,B). The IPSCs in the studied projections were GABAergic, as they were blocked by the GABA_A_ receptor antagonist bicuculine, and disynaptic in nature, as they were blocked by the AMPA/kainate receptor antagonist NBQX (Fig. 5A).

**Figure 5.**
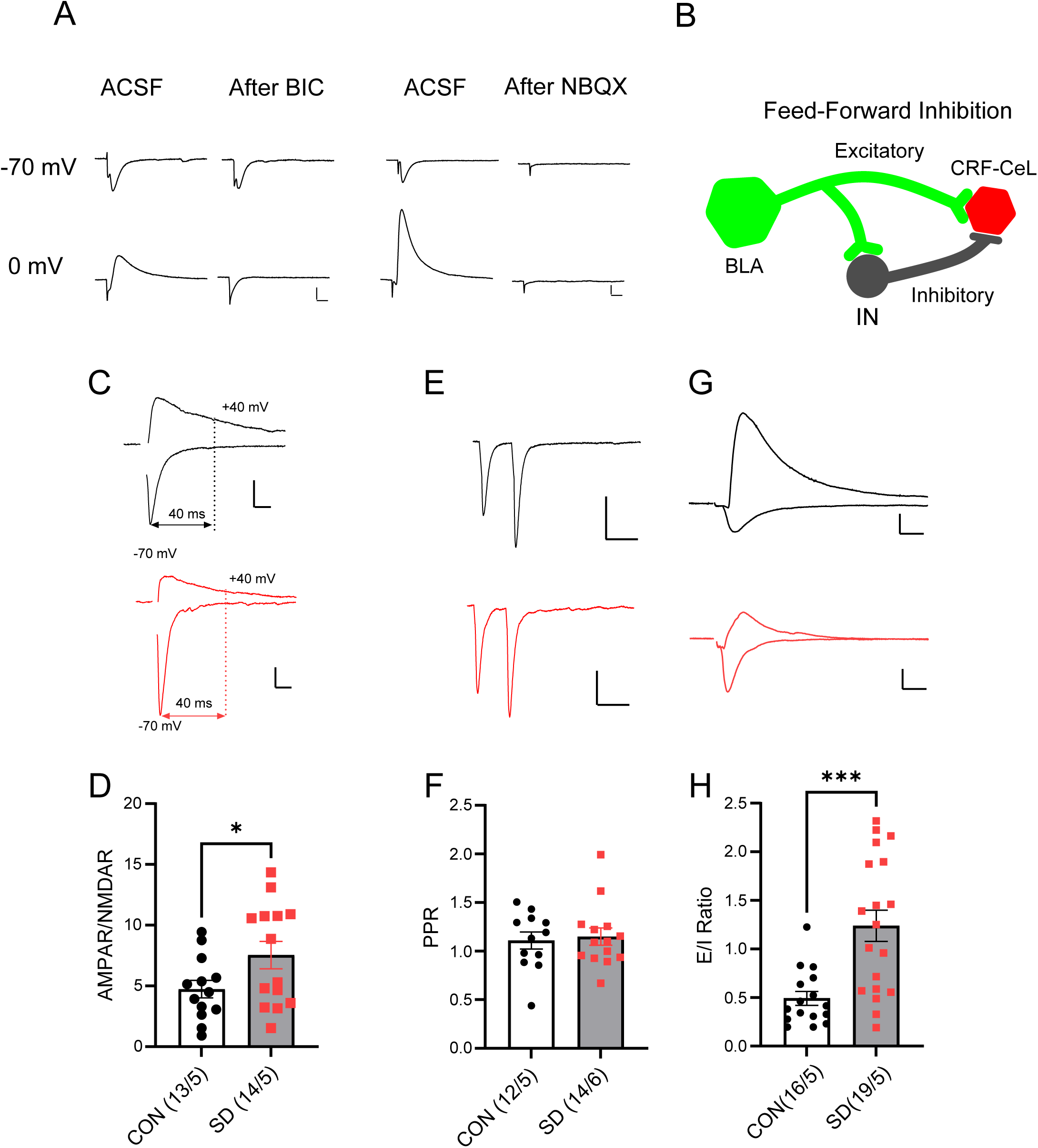
The efficacy of synaptic transmission in BLA-CRF^CeL^ projections was enhanced after SD in female mice. **A**, GABA_A_ receptor antagonist (bicuculline, BIC) blocked inhibitory synaptic responses in BLA-CRF^CeL^ projections recorded at 0 mV, whereas it had no effect on EPSCs recorded at -70 mV (left). Glutamatergic antagonist NBQX (20 μM) blocked both EPSC and IPSC at BLA inputs to CRF^CeL^ cells (right). Scale bars: 50 pA, 10 ms. **B**, The diagram depicting the local circuit mechanism of feed-forward GABAergic inhibition. **C**, Representative traces of EPSCs in the BLA-CRF^CeL^ projections recorded at -70 mV or +40 mV. The NMDAR mediated current was measured at a holding potential of 40 mV at 40 ms after the peak of the AMPAR EPSC mV at -70. Scale bars: 50 pA, 10 ms. **D**, SD resulted in enhanced AMPAR/NMDAR EPSC amplitude ratio in; CON, 13 neurons from 5 mice; SD, 14 neurons from 5 mice. **E**, Representative traces of paired-pulse BLA-CRF^CeL^ EPSCs in slices from CON (top, black) and SD (lower, red) mice. The inter-stimulus interval was 50 ms. Scale bars: 50 pA, 50 ms. **F**, There was no difference in PPR magnitude in the studied pathway; CON, 12 neurons from 5 mice; SD, 14 neurons from 6 mice. **G**, Representative traces of EPSCs (recorded at -70 mV) and IPSCs (recorded at 0 mV) in the same CRF^CeL^ neurons to estimate the E/I ratio in the studied pathway in CON (top, black) and SD (lower, red) mice. Scale bars: 100 pA, 10 ms. **H**, SD was associated with increased E/I ratio in BLA-CRF^CeL^ projections, CON, 16 neurons from 5 mice; SD, 19 neurons from 5 mice.

As CRF^CeL^ neurons receive glutamatergic inputs from the BLA and play a role in both anxiogenesis and the mechanisms of sleep (Fadda and Fratta, 1997; Pawlyk et al., 2006; Sanford et al., 2017; McCullough et al., 2018, 2020; Pomrenze et al., 2019b, 2019a; Jo et al., 2020), we addressed a possibility that SD might be associated with lasting potentiation of glutamatergic synaptic transmission in BLA projections to CRF^CeL^ neurons. To test whether sleep disruption had an effect on synaptic efficacy in the studied pathway, we compared the AMPAR and NMDAR EPSC amplitude ratio between control and SD mice by recording EPSCs at a holding potential of -70 mV first (for AMPA receptor EPSC that was measured at the peak of synaptic current) and then at +40 mV (for NMDA receptor EPSC that was measured at 40 ms after the peak) in the presence of the GABA_A_ receptor antagonist bicuculine. We found that the AMPAR/NMDAR EPSC amplitude ratio was enhanced in slices from sleep-disrupted mice (unpaired *t*-test, *t*_(25)_ = 2.065, *p* = 0.049) (Fig. 5C,D). On the other hand, paired-pulse ratio (PPR; a measure of presynaptic function; Cho et al., 2013) of the EPSCs induced by two closely spaced presynaptic stimuli (50-ms inter-stimulus interval) remained unchanged (unpaired *t*-test, *t_(_*_24)_ = 0.321, *p* = 0.751; Fig. 5 E,F). These findings indicate that sleep disruption-associated synaptic enhancements in projections from the BLA to CRF^CeL^ neurons could be postsynaptically expressed.

As BLA projections to the CeL trigger feed-forward GABAergic inhibition in CRF^CeL^ neurons (Fig. 5A,B), we assessed the effect of sleep disruption on the excitation/inhibition (E/I) ratio in the BLA to CRF^CeL^ inputs by recording monosynaptic glutamatergic EPCSs and disynaptic GABAergic IPSCs in the same neurons, at holding potentials of -70 mV or 0 mV, respectively. We found that the E/I ratio was increased in sleep-disrupted mice (unpaired *t*-test, *t*_(33)_ = 3.973, *p* = 0.0004) (Fig. 5G,H), suggesting that the functional efficiency of excitation in BLA inputs to CRF^CeL^ cells may be increased following sleep disruption.

## Discussion

Our findings provide evidence that the delivery of a single episode of sleep disruption during the light (non-active) phase of the light:dark cycle may be associated with enhanced anxiety-like behaviors and lasting synaptic potentiation in projections from the BLA to CRF^CeL^ neurons in female mice. The observed synaptic strengthening in the studied projections was postsynaptically expressed, as evidenced by the increased AMPAR/NMDAR EPSC amplitude ratio, and it was not accompanied by the changes in intrinsic membrane properties, including neuronal excitability, in SD mice. The SD-associated LTP-like synaptic enhancements in glutamatergic BLA projections to CRF^CeL^ neurons resulted in a shift between excitation and inhibition in this pathway towards a greater functional efficiency of excitation, possibly enhancing synaptically-driven spike output of CRF positive neurons, and thus modifying the signal flow in the anxiety-controlling BLA-CRF^CeL^ circuits, contributing to anxiety enhancements in SD mice. Whereas the effect of SD in human subjects was found to be consistently anxiogenic (Pires et al., 2016a), previous animal studies reported contradictory results regarding the effects of sleep disruption on anxiety levels, with some studies indicating that SD might be anxiogenic, and other groups suggesting that SD may have anxiolytic effects (Andersen et al., 2005; Kumar and Garg, 2008; Cortese et al., 2010; Vollert et al., 2011). The reason for these inconsistencies in reported findings is presently unclear. However several factors could contribute to the experimental variability, including the different species used, duration of sleep disruption, phase of the light:dark cycle being deprived, and specific methods which were used to probe the effects of SD on anxiety-like behaviors (Pires et al., 2013, 2015, 2016b). Notably, in addition to anxiety enhancements, sleep disruption could also trigger mania like behaviors in both human and rodent subjects (Wehr, 1992; Gessa et al., 1995; Wu et al., 2023), which could contaminate the anxiety readouts tested by EPM or OF tests in animal studies. However, we did not detect any changes in the total distance traveled in EPM or OF in SD mice, and, thus, there was no SD-triggered mania-like hyperactivity in our experiments. We focused in our analysis on female mice considering that women are more susceptible to sleep disruption-induced anxiety than men (Goldstein-Piekarski et al., 2018). It is still unclear whether sex differences could be observed in respect to the effects of SD on anxiety levels and/or neurotransmission in BLA-CRF^CeL^ circuits, which we will address in future studies.

The logistically determined limitation of our experimental approach is that mice from CON and SD groups were tested at different periods of time after the 6 hour-long sleep fragmentation chamber exposure. Thus, the testing of mice in SD group has started immediately after the treatment, which took place during an inactive phase of the light:dark cycle from 7:00 AM to 1:00 PM, whereas the testing of CON mice (treated for 6 hours from 7:00 PM to 1:00 AM in their awake cycle) was delayed until 1:00 PM (the same time point as for SD group). To resolve the uncertainties with data interpretation, it would be necessary to analyze two additional control groups, including testing the mice in CON group immediately after the exposure to the sleep fragmentation chamber at 1:00 AM, and testing undisturbed mice which will not be exposed to the sleep fragmentation procedures.

It has been demonstrated previously that CRF^CeL^ neurons may contribute to control of anxogenesis, as activation of these cells promotes anxiety-like behaviors, possibly through CRF and/or dynorphin release (Pomrenze et al., 2019a, 2019b). BLA, especially its posterior parts, provides strong monosynaptic glutamatergic inputs to CRF^CeL^ neurons (Fadok et al., 2017; Fig. 5 in this study), and is involved in anxiety mechanisms (Wang et al., 2011; Tye et al., 2011; Felix-Ortiz et al., 2013). Notably, the CeL contains two distinct subpopulations of GABAergic neurons, termed CeL_on_/PKC-δ^-^ (fear-on) and CeL_off_/PKC-δ^+^ (fear-off) neurons (Ciocchi et al., 2010; Haubensak et al., 2010), with CRF positive cells constituting a fraction of PKC-δ^-^ cells. Moreover, we found previously that axonal fibers containing pituitary adenylate-cyclase-activating polypeptide (PACAP) project to the CeL (Cho et al., 2012; Meloni et al, 2024), and that exogenously applied PACAP potentiates glutamatergic synaptic transmission in BLA-CeL projections, increasing their synaptically triggered spike output (Cho et al., 2012). It is possible that SD could contribute to anxiety enhancements through the induction of PACAP-dependent long-term potentiation of synaptic transmission at BLA inputs to CRF^CeL^ cells specifically. This SD-triggered potentiation would lead to inhibition of CeL_off_/PKC-δ^+^ neurons, disinhibiting CeM and resulting in enhanced fear. Thus, our findings suggest that sleep disruption may enhance anxiety levels through the induction of SD-associated synaptic plasticity in BLA-CeL circuits.

## Abbreviations

BLA: basolateral amygdala
CeA: central nucleus of amygdala
CeL: lateral subnucleus of CeA
CRF: corticotropin-releasing factor
EPSCs: excitatory postsynaptic currents
IPSCs: inhibitory postsynaptic currents
4-AP: 4-aminopyridine
AMPAR: α-amino-3-hydroxy-5-methyl-4-isoxazolepropionic acid receptor
NMDAR: N-methyl-D-aspartate receptor
GABA: γ-Aminobutyric acid
SD: sleep disruption

## Acknowledgements

This work was supported by grants P50MH115874 (to KJR and WAC), R01MH108665 (to KJR and VYB), and R01MH123993 (to V.Y.B).

## References

Ahrens, S., Wu, M.V., Furlan, A., Hwang, G.R., Paik, R., Li, H., Penzo, M.A., Tollkuhn, J. & Li, B. (2018) A central extended amygdala circuit that modulates anxiety. J. Neurosci. 38, 5567–5583.

Altemus, M., Sarvaiya, N. & Epperson, C.N. (2014) Sex differences in anxiety and depression clinical perspectives. Front. Neuroendocrinol. 35, 320–330.

Alvaro, P.K., Roberts, R.M. & Harris, J.K. (2013) A systematic review assessing bidirectionality between sleep disturbances, anxiety, and depression. Sleep 36, 1059–1068.

Andersen, M.L., Perry, J.C. & Tufik, S. (2005) Acute cocaine effects in paradoxical sleep deprived male rats. Prog. Neuropsychopharmacol. Biol. Psychiatry 29, 245–251.

Babson, K.A., Trainor, C.D., Feldner, M.T. & Blumenthal, H. (2010) A test of the effects of acute sleep deprivation on general and specific self-reported anxiety and depressive symptoms: An experimental extension. J. Behav. Ther. Exp. Psychiatry 41, 297–303.

Baimel, C., Lau, B.K., Qiao, M. & Borgland, S.L. (2017) Projection-Target-Defined Effects of Orexin and Dynorphin on VTA Dopamine Neurons. Cell Rep. 18, 1346–1355.

Besedovsky, L., Lange, T. & Haack, M. (2019) The sleep-immune crosstalk in health and disease. Physiol. Rev. 99, 1325–1380.

Bian, W.J., Brewer, C.L., Kauer, J.A. & De Lecea, L. (2022) Adolescent sleep shapes social novelty preference in mice. Nat. Neurosci. 25, 912–923.

Bourin, M., Dhonnchadha, B.Á.N., Colombel, M.C., Dib, M. & Hascoët, M. (2001) Cyameazine as an anxiolytic drug on the elevated plus maze and light/dark paradigm in mice. Behavioural Brain Research, 124, 87–95.

Brownstone, R.M., Jordan, L.M., Kriellaars, D.J., Noga, B.R. & Shefchyk, S.J., (1992) On the regulation of repetitive firing in lumbar motoneurones during fictive locomotion in the cat. Exp. Brain Res. 90, 441–455.

Cai, H., Haubensak, W., Anthony, T.E. & Anderson, D.J. (2014) Central amygdala PKC-δ+ neurons mediate the influence of multiple anorexigenic signals. Nat. Neurosci. 17, 1240–1248.

Cho, J.H., Zushida, K., Shumyatsky, G.P., Carlezon, W.A., Meloni, E.G. & Bolshakov, V.Y. (2012) Pituitary adenylate cyclase-activating polypeptide induces postsynaptically expressed potentiation in the intra-amygdala circuit. J. Neurosci. 32, 14165–14177.

Cho, J.H., Deisseroth, K. & Bolshakov, V.Y. (2013) Synaptic encoding of fear extinction in mPFC-amygdala circuits. Neuron 80, 1491–507

Ciocchi, S., Herry, C., Grenier, F., Wolff, S.B.E., Letzkus, J.J., Vlachos, I., Ehrlich, I., Sprengel, R., Deisseroth, K., Stadler, M.B., Müller, C. & Lüthi, A. (2010) Encoding of conditioned fear in central amygdala inhibitory circuits. Nature 468, 277–282.

Cortese, B.M., Mitchell, T.R., Galloway, M.P., Prevost, K.E., Fang, J., Moore, G.J. & Uhde, T.W. (2010) Region-specific alteration in brain glutamate: Possible relationship to risk-taking behavior. Physiol. Behav. 99, 445–450.

Cox, R.C. & Olatunji, B.O. (2016) A systematic review of sleep disturbance in anxiety and related disorders. J. Anxiety Disord. 37, 104–129.

Davis, M. (1992) The role of the amygdala in fear and anxiety. Annu. Rev. Neurosci. 15, 353–375.

Douceau, S., Lemarchand, E., Hommet, Y., Lebouvier, L., Joséphine, C., Bemelmans, A.P., Maubert, E., Agin, V. & Vivien, D. (2022) PKCδ-positive GABAergic neurons in the central amygdala exhibit tissue-type plasminogen activator: role in the control of anxiety. Mol. Psychiatry 27, 2197–2205.

Fadda, P. & Fratta, W. (1997) Stress-induced sleep deprivation modifies corticotropin releasing factor (CRF) levels and CRF binding in rat brain and pituitary. Pharmacol. Res. 35, 443–446.

Fadok, J.P., Krabbe, S., Markovic, M., Courtin, J., Xu, C., Massi, L., Botta, P., Bylund, K., Müller, C., Kovacevic, A., Tovote, P. & Lüthi, A. (2017) A competitive inhibitory circuit for selection of active and passive fear responses. Nature 542, 96–100.

Felix-Ortiz, A.C., Beyeler, A., Seo, C., Leppla, C.A., Wildes, C.P. & Tye, K.M. (2013) BLA to vHPC inputs modulate anxiety-related behaviors. Neuron 79, 658–664.

Foilb, A.R., Taylor-Yeremeeva, E.M., Fritsch, E.L., Ravichandran, C., Lezak, K.R., Missig, G., McCullough, K.M. & Carlezon, W.A. (2023) Differential effects of the stress peptides PACAP and CRF on sleep architecture in mice. bioRxiv 2023.03.22.533872.

Foilb, A.R., Taylor-Yeremeeva, E.M., Schmidt, B.D., Ressler, K.J. & Carlezon, W.A. (2024) Acute sleep deprivation reduces fear memories in male and female mice. bioRxiv 10.1101/2024.01.30.577985

Gessa, G.L., Pani, L., Fadda, P. & Fratta, W. (1995) Sleep deprivation in the rat: an animal model of mania. Eur. Neuropsychopharmacol. 5, 89–93.

Goldstein-Piekarski, A.N., Greer, S.M., Saletin, J.M., Harvey, A.G., Williams, L.M. & Walker, M.P. (2018) Sex, sleep deprivation, and the anxious brain. J. Cogn. Neurosci. 30, 565–578.

Hajali, V., Sheibani, V., Esmaeili-Mahani, S. & Shabani, M. (2012) Female rats are more susceptible to the deleterious effects of paradoxical sleep deprivation on cognitive performance. Behav. Brain Res. 228, 311–318.

Hasegawa, E., Miyasaka, A., Sakurai, K., Cherasse, Y., Li, Y. & Sakurai, T. (2022) Rapid eye movement sleep is initiated by basolateral amygdala dopamine signaling in mice. Science 375, 994–1000.

Haubensak, W., Kunwar, P.S., Cai, H., Ciocchi, S., Wall, N.R., Ponnusamy, R., Biag, J., Dong, H.-W., Deisseroth, K., Callaway, E.M., Fanselow, M.S., Lüthi, A. & Anderson, D.J. (2010) Genetic dissection of an amygdala microcircuit that gates conditioned fear. Nature 468, 270–276.

Hillman, D.R. & Lack, L.C. (2013) Public health implications of sleep loss: the community burden. Med. J. Aust. 199, S7–10.

Jo, Y.S., Namboodiri, V.M.K., Stuber, G.D. & Zweifel, L.S. (2020) Persistent activation of central amygdala CRF neurons helps drive the immediate fear extinction deficit. Nat. Commun. 11:422.

Johnson, E.O., Roth, T. & Breslau, N. (2006) The association of insomnia with anxiety disorders and depression: Exploration of the direction of risk. J. Psychiatr. Res. 40, 700–708.

Kessler, R.C., McGonagle, K.A., Zhao, S., Nelson, C.B., Hughes, M., Eshleman, S., Wittchen, H.U. & Kendler, K.S. (1994) Lifetime and 12-month prevalence of DSM-III-R psychiatric disorders in the United States: Results from the national comorbidity survey. Arch. Gen. Psychiatry 51, 8–19.

Khan, M.S. & Aouad, R. (2022) The Effects of insomnia and sleep loss on cardiovascular disease. Sleep Med. Clin. 17, 193–203.

Kim, S.Y., Adhikari, A., Lee, S.Y., Marshel, J.H., Kim, C.K., Mallory, C.S., Lo, M., Pak, S., Mattis, J., Lim, B.K., Malenka, R.C., Warden, M.R., Neve, R., Tye, K.M. & Deisseroth K. (2013). Diverging neural pathways assemble a behavioural state from separable features in anxiety. Nature 496, 219–223.

Koehl, M., Battle, S.E. & Turek, F.W. (2003) Sleep in female mice: a strain comparison across the estrous cycle. Sleep 26, 267–272.

Kumar, A. & Garg, R. (2008) A role of nitric oxide mechanism involved in the protective effects of venlafaxine in sleep deprivation. Behav. Brain Res. 194, 169–173.

Ma, C., Zhong, P., Liu, D., Barger, Z.K., Zhou, L., Chang, W.C., Kim, B. & Dan, Y. (2019) Sleep regulation by neurotensinergic neurons in a thalamo-amygdala circuit. Neuron 103, 323–334.

Madan Jha V. (2023) The prevalence of sleep loss and sleep disorders in young and old adults. Aging Brain 3:100057.

McCullough, K.M., Chatzinakos, C., Hartmann, J., Missig, G., Neve, R.L., Fenster, R.J., Carlezon, W.A., Daskalakis, N.P. & Ressler, K.J. (2020) Genome-wide translational profiling of amygdala Crh-expressing neurons reveals role for CREB in fear extinction learning. Nat. Commun. 11:5180.

McCullough, K.M., Morrison, F.G., Hartmann, J., Carlezon, W.A. & Ressler, K.J. (2018) Quantified coexpression analysis of central amygdala subpopulations. eNeuro 5(1):ENEURO.0010-18.2018.

Meloni, E.G., Carlezon WA Jr & Bolshakov V.Y. (2024) Association between social dominance hierarchy and PACAP expression in the extended amygdala, corticosterone, and behavior in C57BL/6 male mice. Sci Rep. 14(1):8919.

Montes-Rodríguez, C.J., Rueda-Orozco, P.E. & Prospéro-García, O. (2019) Total sleep deprivation impairs fear memory retrieval by decreasing the basolateral amygdala activity. Brain Res. 1719, 17–23.

Nair, D., Zhang, S.X.L., Ramesh, V., Hakim, F., Kaushal, N., Wang, Y. & Gozal, D. (2011) Sleep fragmentation induces cognitive deficits via nicotinamide adenine dinucleotide phosphate oxidase-dependent pathways in mouse. Am. J. Respir. Crit. Care Med. 184, 1305–1312.

Ohayon, M.M. & Roth, T. (2003) Place of chronic insomnia in the course of depressive and anxiety disorders. J. Psychiatr. Res. 37, 9–15.

Palma, J.-A., Urrestarazu, E. & Iriarte, J. (2013) Sleep loss as risk factor for neurologic disorders: a review. Sleep Med. 14, 229–236.

Paul, K.N., Dugovic, C., Turek, F.W. & Laposky, A.D. (2006) Diurnal sex differences in the sleep-wake cycle of mice are dependent on gonadal function. Sleep 29, 1211–1223.

Pawlyk, A., Sanford, L., Brennan, F., Morrison, A. & Ross, R. (2006) Corticotropin-releasing factor microinjection into the central nucleus of the amygdala alters REM sleep. Pharmacol. Rep. 58, 125–130.

Pires, G.N., Bezerra, A.G., Tufik, S. & Andersen, M.L. (2016a) Effects of acute sleep deprivation on state anxiety levels: a systematic review and meta-analysis. Sleep Med. 24, 109–118.

Pires, G.N., Bezerra, A.G., Tufik, S. & Andersen, M.L. (2016b) Effects of experimental sleep deprivation on anxiety-like behavior in animal research: Systematic review and meta-analysis. Neurosci. Biobehav. Rev. 68, 575–589.

Pires, G.N., Tufik, S. & Andersen, M.L. (2015) Sleep deprivation and anxiety in humans and rodents—translational considerations and hypotheses. Behav. Neurosci. 129, 621–633.

Pires, G.N., Tufik, S. & Andersen, M.L. (2013) Grooming analysis algorithm: Use in the relationship between sleep deprivation and anxiety-like behavior. Prog. Neuropsychopharmacol. Biol. Psychiatry 41, 6–10.

Pomrenze, M.B., Giovanetti, S.M., Maiya, R., Gordon, A.G., Kreeger, L.J. & Messing, R.O. (2019a) Dissecting the roles of GABA and neuropeptides from rat central amygdala CRF neurons in anxiety and fear learning. Cell Rep. 29, 13–21.

Pomrenze, M.B., Tovar-Diaz, J., Blasio, A., Maiya, R., Giovanetti, S.M., Lei, K., Morikawa, H., Hopf, F.W. & Messing, R.O. (2019b) A corticotropin releasing factor network in the extended amygdala for anxiety. J. Neurosci. 39, 1030–1043.

Ramsawh, H.J., Stein, M.B., Belik, S.L., Jacobi, F. & Sareen, J. (2009) Relationship of anxiety disorders, sleep quality, and functional impairment in a community sample. J. Psychiatr. Res. 43, 926–933.

Riccio, A., Li, Y., Moon, J., Kim, K.S., Smith, K.S., Rudolph, U., Gapon, S., Yao, G.L., Tsvetkov, E., Rodig, S.J., Van’t Veer, A., Meloni, E.G., Carlezon, W.A., Bolshakov, V.Y. & Clapham D.E. (2009). Essential role for TRPC5 in amygdala function and fear-related behavior. Cell 137, 761–772.

Royer, S., Martina, M. & Paré, D. (1999) An inhibitory interface gates impulse traffic between the input and output stations of the amygdala. J. Neurosci. 19, 10575–10583.

Sah, P., Faber, E.S.L., Lopez De Armentia, M. & Power, J. (2003) The Amygdaloid Complex: Anatomy and Physiology. Physiol. Rev. 83, 803–834.

Sanders, S.K. & Shekhar, A. (1995) Regulation of anxiety by GABAA receptors in the rat amygdala. Pharmacol. Biochem. Behav. 52, 701–706.

Sanford, C.A., Soden, M.E., Baird, M.A., Miller, S.M., Schulkin, J., Palmiter, R.D., Clark, M. & Zweifel, L.S. (2017) A central amygdala CRF circuit facilitates learning about weak threats. Neuron 93, 164–178.

Selvi, Y., Gulec, M., Agargun, M.Y. & Besiroglu, L. (2007) Mood changes after sleep deprivation in morningness–eveningness chronotypes in healthy individuals. J. Sleep Res. 16, 241–244.

Shin, L.M., Rauch, S.L. & Pitman, R.K. (2006) Amygdala, medial prefrontal cortex, and hippocampal function in PTSD. Ann. N. Y. Acad. Sci. 1071, 67–79.

Sun, Y., Qian, L., Xu, L., Hunt, S. & Sah, P. (2023) Somatostatin neurons in the central amygdala mediate anxiety by disinhibition of the central sublenticular extended amygdala. Mol. Psychiatry 28, 4163–4174.

Tye, K.M., Prakash, R., Kim, S.Y., Fenno, L.E., Grosenick, L., Zarabi, H., Thompson, K.R., Gradinaru, V., Ramakrishnan, C. & Deisseroth, K. (2011) Amygdala circuitry mediating reversible and bidirectional control of anxiety. Nature 471, 358–362.

van der Helm, E., Yao, J., Dutt, S., Rao, V., Saletin, J.M. & Walker, M.P. (2011) REM sleep depotentiates amygdala activity to previous emotional experiences. Curr. Biol. 21, 2029–2032.

Vollert, C., Zagaar, M., Hovatta, I., Taneja, M., Vu, A., Dao, A., Levine, A., Alkadhi, K. & Salim, S. (2011) Exercise prevents sleep deprivation-associated anxiety-like behavior in rats: potential role of oxidative stress mechanisms. Behav. Brain Res. 224, 233–240.

Wang, D.V., Wang, F., Liu, J., Zhang, L., Wang, Z. & Lin, L. (2011) Neurons in the amygdala with response-selectivity for anxiety in two ethologically based tests. PLoS ONE 6(4): e18739

Wang, Y., Liu, Z., Cai, L., Guo, R., Dong, Y. & Huang, Y.H. (2020) A critical role of basolateral amygdala–to–nucleus accumbens projection in sleep regulation of reward seeking. Biol. Psychiatry 87, 954–966.

Wehr, T.A. (1992) Improvement of depression and triggering of mania by sleep deprivation. JAMA 267, 548–551.

Wu, M., Zhang, X., Feng, S., Freda, S.N., Kumari, P., Dumrongprechachan, V. & Kozorovitskiy, Y. (2023) Dopamine pathways mediating affective state transitions after sleep loss. Neuron 112, 141–154

Yoo, S.-S., Gujar, N., Hu, P., Jolesz, F.A. & Walker, M.P. (2007) The human emotional brain without sleep — a prefrontal amygdala disconnect. Curr. Biol. 17, R877–R878.

Zhang, B., Wing, Y.K., 2006. Sex differences in insomnia: a meta-analysis. Sleep 29, 85–93.

